# TRPV4 Inhibition by a Natural Product Mediates Analgesia

**DOI:** 10.64898/2026.01.05.697673

**Authors:** Zhen Yuan, Jie Qiu, Shiliang Li, Hanqing Zhao, Na Chen, Shanshan Ruan, Yu Gan, Jiawen Wang, Tianyu Ye, Wenjie Liao, Ziyuan Li, Yanyan Diao, Lili Zhu, Zhenjiang Zhao, Rui Wang, Yongjun Zheng, Jin Liu, Honglin Li, Bowen Ke

## Abstract

The transient receptor potential vanilloid 4 (TRPV4) channel, a polymodal calcium-permeable cation channel, is activated by diverse stimuli, including hyposmolarity, moderate heat, mechanical stress, and acidic pH. TRPV4 plays a critical role in the onset and progression of pain and is implicated in multiple pathological conditions, positioning it as an attractive therapeutic target. However, the development of TRPV4-targeted therapeutics has been hindered by limited mechanistic understanding of its inhibition. Here, the natural product (*R*)-1-(3-ethylphenyl)ethane-1,2-diol (AH001) is identified as a potent, subtype-selective TRPV4 inhibitor. A potential gating mechanism of TRPV4 inhibition is proposed through a combination of structural analyses and site-directed mutagenesis. Electrophysiological studies and cryo-electron microscopy (cryo-EM) further reveal that AH001’s glucuronide metabolite, AHP, retains robust TRPV4 inhibitory activity, providing a critical structural basis for rational optimization of AH001 as a lead compound. Notably, AH001 demonstrated significant efficacy in reducing neuronal hyperexcitability in dorsal root ganglia (DRGs) and produced pronounced analgesic effects across multiple pain models, highlighting its therapeutic potential in pain relief. This study offers critical structural and mechanistic insights into TRPV4 inhibition and lays the groundwork for the structure-based design of TRPV4-targeted therapeutics.

## 1. Introduction

Pain serves as a fundamental physiological warning system essential for the survival of living organisms^1, 2^. However, chronic pain is a multidimensional and subjective condition that profoundly impairs the physical and psychological well-being of millions of individuals worldwide^3, 4^, representing a major unmet medical need due to the limited efficacy and significant side effects of current treatments. Among the molecular mediators involved in nociception, the transient receptor potential (TRP) channel family has garnered considerable attention for its unique electrophysiological responses to diverse stimuli and chemical stimuli^5, 6^.

Within this family, transient receptor potential vanilloid 4 (TRPV4), a polymodal and non-selective cation channel characterized by high Ca^2+^ permeability^7, 8^, is prominently expressed in various tissues, including the nervous system, respiratory system, kidney, and skin, where it responds to stimuli such as hyperosmolarity, moderate heat, mechanical stress, and acidic pH, underscoring its versatile physiological roles in calcium signaling, thermosensation, cellular volume maintenance, and energy homeostasis^9, 10^. Dysregulation of TRPV4 has been implicated in the pathogenesis of multiple conditions, including chronic pain, osteoarthritis, neuropathic pain, and neurodegenerative disorders, establishing it as a highly compelling therapeutic target^11–13^.

Despite its therapeutic promise, the development of TRPV4-specific inhibitors has been constrained by several challenges, notably the high structural and sequence homology within the TRPV subfamily, which complicates the design of subtype-selective modulators^14^. Recent advances in structural biology, approaches, particularly cryo-electron microscopy (cryo-EM), have significantly enhanced our understanding of TRPV4 activation dynamics by elucidating its conformational states. However, the detailed mechanisms by which TRPV4 antagonists exert their effects, particularly via allosteric modulation of channel gating, remain incompletely characterized. Recent studies^15–18^ on TRPV4 antagonists (e.g., GSK2798745, HC-067047, GSK3527497) and agonists (GSK1016790A, 4α-PDD, agonist-1) have shed light on how they bind to the voltage sensor-like domain (VSLD) and the TRP domain to modulate channel activity. However, the detailed structural and mechanistic basis of allosteric regulation of TRPV4 remains elusive, posing a significant barrier to the development of potent and selective inhibitors.

In our previous study, the natural product AH001, discovered through virtual screening, inhibited GSK1016790A-stimulated TRPV4 channel-mediated calcium influx, and we resolved the cryo-EM structure of AH001 bound to TRPV4 (PDB ID: 9IQX)^19^. In this study, whole-cell patch-clamp recordings further demonstrate that AH001 serves as a potent and selective antagonist of TRPV4 channels. Detailed structural analysis, including single-point/multipoint mutagenesis experiments, revealed the potential gating mechanism by which this selective inhibitor modulates TRPV4 function. Our findings show that AH001 and its glucuronide metabolite, AHP, effectively bind to TRPV4 and induce a conformational change that blocks channel activity. Furthermore, we provided robust evidence of AH001’s analgesic activity *in vitro* and *in vivo* through TRPV4 inhibition. Collectively, our cryo-EM structure determination, mechanistic investigation, and functional validation not only provide novel molecular insights into the molecular mechanisms underlying TRPV4 inhibition but also lay a robust foundation for the structure-guided design of targeted analgesics.

## 2. Results

### 2.1 Functional characterization of TRPV4 inhibition by AH001 and its active metabolite AHP

We expressed human TRPV4 (hTRPV4) in HEK 293 cells and assessed its functionality using whole-cell patch-clamp recordings. Sustained exposure to GSK1016790A (abbreviated GSK101) led to a progressive enhancement in current amplitude, followed by stabilization at a plateau that persisted for approximately 10 minutes. This stability confirms the robustness and reliability of our whole-cell recordings assay (**Figure 1a**). Subsequently, a high-throughput virtual screening identified the natural product (*R*)-1-(3-ethylphenyl)ethane-1,2-diol^20^ (named AH001) as a potent inhibitor of human TRPV4 (**Figure 1b-c**). AH001 exhibited a concentration-dependent reduction of TRPV4 currents (**Figure 1d**), with a half-maximum inhibitory concentration (IC_50_) of 2.27 μM (**Figure 1e**). Pharmacokinetic analyses revealed that AH001 undergoes primarily metabolism *in vivo* via phase II pathways, yielding a glucuronide conjugate metabolite (named AHP, **Figure 1c**; **Figure S1**, Supporting Information). Remarkably, this metabolite retained inhibitory active against TRPV4, exhibiting dose-dependent inhibition (**Figure 1f**) with an IC_50_ of 6.10 μM (**Figure 1g**).

**Figure 1.**
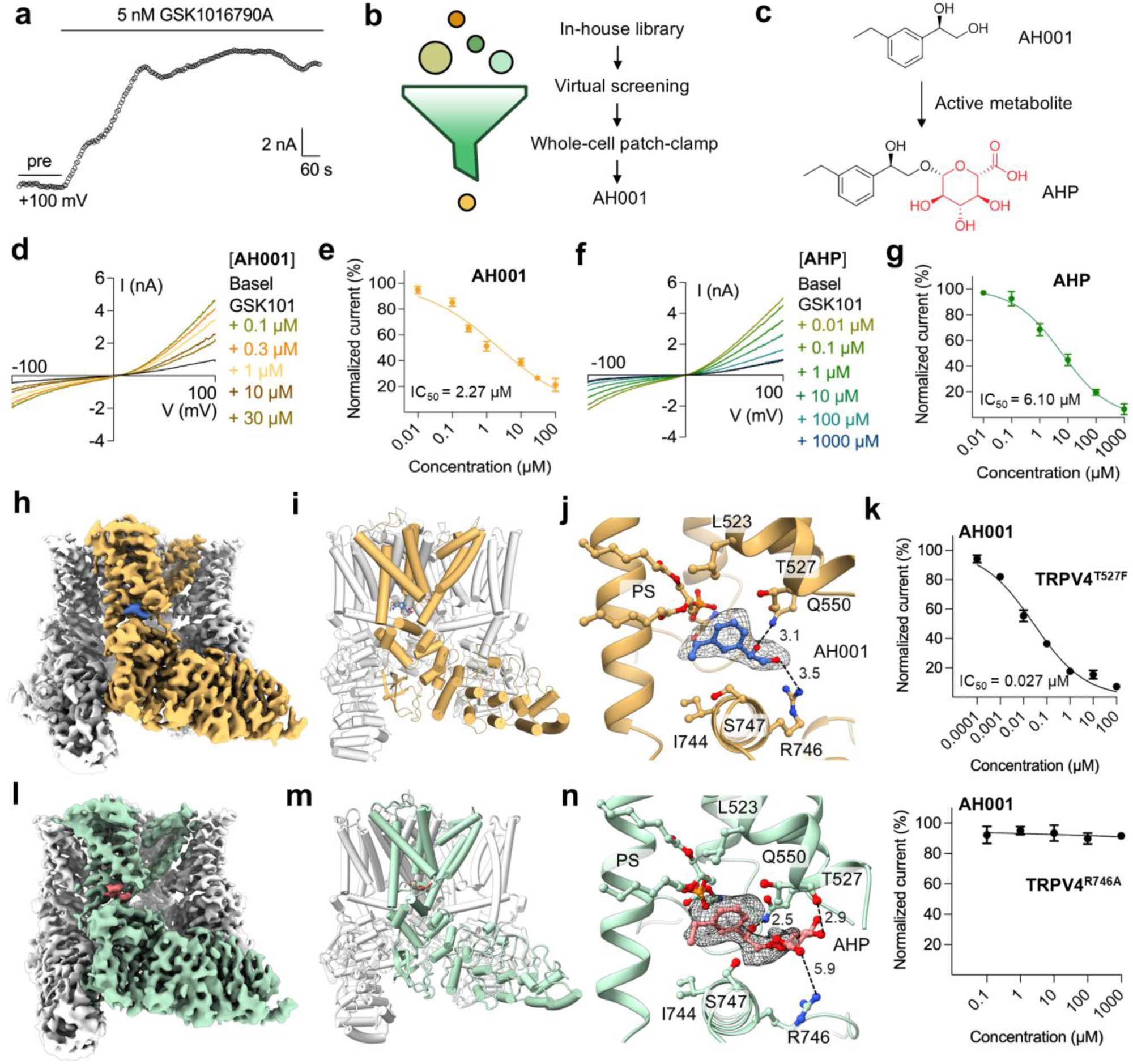
Identification of AH001 and its active metabolite AHP as TRPV4 inhibitors. **a**, The time course at +100 mV showing the effect of GSK101 on hTRPV4 expressed in HEK293 cells. **b**, Virtual screening workflow for the discovery of TRPV4 inhibitor. **c**, Chemical structure of AH001 and its glucuronide metabolite AHP. **d and f**, *I*-*V* curves of currents of keratinocytes inhibited by AH001 (**d**) or AHP (**f**) in the presence of 5 nM GSK101. **e and g**, Dose-response analysis of different concentrations of AH001 (**f)** or AHP (**i)** on GSK101-evoked currents of hTRPV4 at +100 mV. **h and l**, Cryo-EM structures of hTRPV4 in the AH001-bound (**h**) and AHP-bound states (**l**). EM densities of AH001 and AHP are represented as blue and red surfaces, respectively. **i and m**, The partial structure of AH001 bound to TRPV4 viewed parallel to the membrane (**i**). TRPV4_AHP_ structure viewed parallel to the membrane (**m**).**j and n**, Zoomed-in view of the AH001- and AHP-binding pocket. AH001 and AHP are represented as blue and red sticks, respectively. Residues close to AH001 or AHP are shown in stick representation. EM density for AH001 and AHP is contoured at 5σ (gray mesh). The black dashed lines represent the distances (in Å) between heavy atoms involved in hydrogen bonds. **k**, Representative time course and curve fitting of dose-dependent inhition of ramp current at +100 mV by AH001 on hTRPV4-T527F and hTRPV4-R746A mutants.

Building upon our prior structural determination of the AH001 bound to TRPV4 (PDB ID: 9IQX, **Figure 1h-i**), we extended cryo-EM single-particle analysis to its glucuronide metabolite AHP (**Figure S2-4**; **Table S1**, Supporting Information). Following heterogeneous refinement, nonuniform refinement with two-fold rotational symmetry (C2) was applied to capture the primary conformational state, resulting in a high-quality 3D map of TRPV4 at a global resolution of 3.48 Å (**Figure 1I**; **Figure S3**, Supporting Information). This density map enabled the precise reconstruction of TRPV4 3D structure (**Figure 1m**). We identified a distinct non-protein density within a triangle-shaped cavity (named VSLD cavity) formed by S1-S4 bundles and TRP helix (**Figure 1n**). Notably, this density was larger than that observed for AH001 (PDB ID: 9IQX) and overlapped with AH001’s density, confirming it as the binding site of the AHP (**Figure 1n**; **Figure S5a**, Supporting Information). Importantly, non-protein electron densities observed within the VSLD cavity could be reasonably accommodated to the corresponding antagonists, particularly AH001, thereby revealing the binding sites of the two antagonists (**Figure 1j, n**). In the case of AH001, R746 and Q550 form hydrogen bonds with the hydroxyl tail of the ligand, while T527, L523, and I744 mediate hydrophobic interactions with the aromatic head of the ligand (**Figure 1j**). Notably, the short side chain residue T527 plays crucial role in making space in the cleft to accommodate the head of AH001 (**Figure 1j**). To evaluate the role of T527 in AH001 binding, we introduced the T527F mutation, hypothesizing that the aromatic ring of phenylalanine could engage in π-π stacking interaction with the ligand, thereby enhancing its potency. As expected, the T527F mutation increased AH001 potency by 84-fold compared to the TRPV4^WT^ (**Figure 1k**), identifying T527 as a pivotal residue for ligand binding. Conversely, the R746A mutation almost completely abolished AH001 potency (**Figure 1k**), indicating the critical contribution of the hydrogen bond formed by R746 to AH001 binding. These data support the observed binding pocket of AH001 and the key interactions. As a metabolite of AH001, compound AHP exhibits a similar binding mode within the pocket. However, due to the higher flexibility of its glucuronide moiety, it likely loses the interaction with R746 while gaining an additional hydrogen bond with the backbone carbonyl group of T527 (**Figure 1n**). The C2-symmetric TRPV4_AHP_ structure reveals that the side chains of residues M718 and V722 formed a hydrophobic seal of the pore, indicating that the inner gate is nonconductive (**Figure S5b-c**, Supporting Information). Therefore, our results suggest that both AH001 and its active metabolite, AHP, bind to the VSLD cavity to inhibit TRPV4 activity, providing mechanistic insights into their inhibitory effects.

### 2.2 AH001 selectively inhibits TRPV4

To assess the subtype selectivity of AH001, we evaluated its inhibitory effects on pain-related channels, including TRPV1, TRPV3, TRPA1, and TRPM8, transiently or stably expressed in HEK293 cells (Figure 2a-d). At a concentration of 100 μM, AH001 inhibited TRPV4 activity by 78.77 ± 5.12%, while demonstrating substantially lower inhibitory effects of TRPV1, TRPV3, TRPA1 and TRPM8, with inhibition rates of 23.84 ± 4.15%, 4.86 ± 1.74%, 2.94 ± 1.42% and 16.76 ± 4.65%, respectively (Figure 2e-f). These findings establish AH001 as a selective inhibitor of TRPV4 over other TRP channels. To explore the molecular determinants of AH001 specificity, we compared the sequence alignments of five TRP channels in the vicinity of the binding pocket (Figure 2g). The sequence alignments revealed significant non-conservation among the residues involved in AH001 interactions, particularly T527, which in TRPV4 is a threonine but is substituted with bulky and charged residues in other TRP channels (Figure 2g). Therefore, we speculate that T527 may serve as a critical residue for the activity and selectivity of AH001.

**Figure 2.**
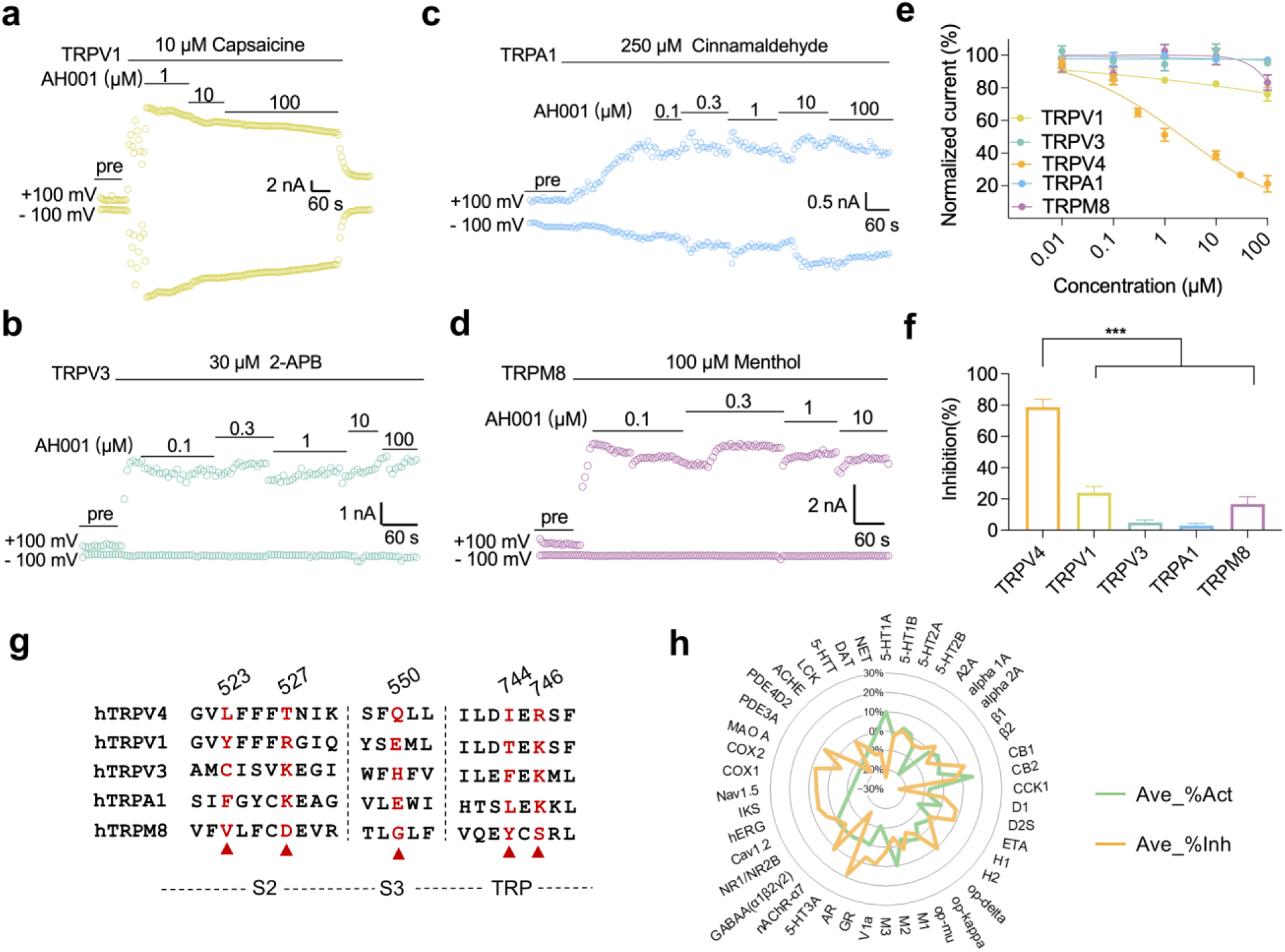
Selectivity evaluation of AH001 on TRPV1, TRPV3, TRPA1, and TRPM8 channels expressed in HEK293 cells. **a-d**, Representative time course of whole-cell TRPV1 (**a**), TRPV3 (**b**), TRPA1 (**c**) and TRPM8 (**d**) currents obtained from −100 mV to +100 mV voltage ramp in HEK293 cells perfused with AH001. **e-f,** Dose‒response analysis of different concentrations of AH001 (**e**) and summary of the percent of inhibition by 100 µM AH001 (**f**) on agonist-evoked currents of hTRPV1 (10 µM capsaicin), hTRPV3 (30 µM 2-APB), hTRPA1 (250 µM cinnamaldehyde) and hTRPM8 (100 µM menthol). One-way ANOVA with Tukey’s post hoc test. Data are shown as the means ± SEMs. \*\*\**P* < 0.001. **g**, Sequence alignment of the human TRPV4, TRPV1, TRPV3, TRPA1 and TRPM8 channels. The red triangles indicate the residues at the AH001 binding pocket. T527 is important for AH001 selectivity. **h**, The interference of native ligand binding of a total of 44 targets by 10 μM AH001, the inhibition (or stimulation) was below 20% in all cases.

To evaluate the safety profile of AH001, we conducted *in vitro* pharmacological profiling against a safety screening panel encompassing 44 targets, as recommended by leading pharmaceutical corporations, including AstraZeneca, GlaxoSmithKline, Novartis, and Pfizer^21^. This panel spans a broad range of biological targets, including ion channels, G protein-coupled receptors (GPCRs), enzymes, nuclear receptors, and transporters. Functional assays revealed that 10 μM AH001 exhibited no significant inhibitory or stimulatory effects on any of the 44 targets, with all inhibition or stimulation rates below 20% (Figure 2h; **Table S2**, Supporting Information). These results suggest that AH001 is a potent and subtype-selective inhibitor of TRPV4 with minimal safety concerns.

### 2.3 Proposal of the potential gating mechanism of TRPV4 inhibition

To reveal how AH001 inhibits TRPV4, we compared our AH001-bound closed TRPV4 structure (PDB ID: 9IQX) and the 4a-PDD-bound open TRPV4 (PDB ID: 8T1D). Compared to the open-state structure of TRPV4, AH001 binding induces substantial conformational changes in the transmembrane domain (TMD) of TRPV4 (Figure 3a). Notably, upon AH001 binding, the S2-S3 helices shift upward in the extracellular direction, resulting in an increase of ∼2 Å in the distance between the S2-S3 and TRP helices (Figure 3b). The decoupled motion between the S2-S3 and TRP regions increases the space between the helices, facilitating the insertion of the S4-S5 linker. To assess the impact of the S2-S3 helix, we introduced alanine substitutions of K535 (S2-S3 linker) and D531 (S2-S3 linker), which significantly reduced the inhibitory efficiency of AH001 (Figure 3f).

**Figure 3.**
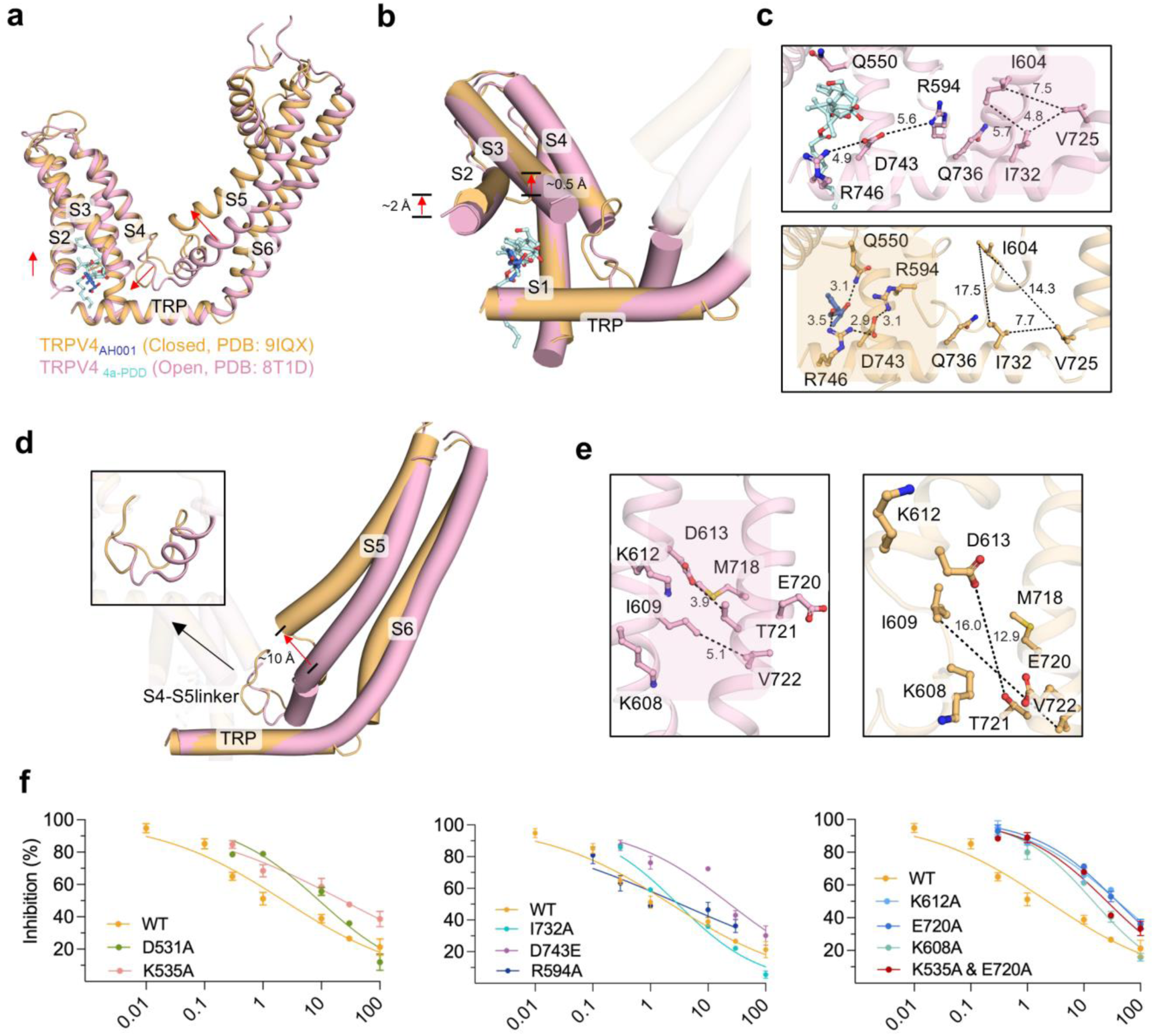
The potential mechanism of TRPV4 inhibition by AH001. **a**, Superposition of the TMDs from TRPV4_AH001_ (closed, yellow) and TRPV4_4α-PDD_ (open, pink). Relative movements of domains are indicated by red arrows. **b**, Side-by-side comparison of the S1-S4 helices and TRP domain rearrangements in the open and closed states. **c**, Comparison of coupling networks at the VSLD, TRP and S4-S5 linker in closed and open states. Dashed lines represent hydrogen bonds, salt bridges, or hydrophobic interactions. The black dashed lines represent the distances (in Å) between heavy atoms. **d**, Side-by-side comparison of the TRP domain and S5-S6 helices rearrangements in the open and closed states. **e**, Comparison of coupling networks at the S5 and S6 in closed and open states. **f**, Curve fitting of dose-dependent inhibition of 5 nM GSK1016790A-evoked currents at +100 mV by AH001 on various TRPV4 mutants compared to WT.

When the AH001 binds to the site (open to closed transition), it engages R594 (S4-S5 linker) and D746 (TRP) into salt bridges with D743 (TRP) (Figure 3b-c). This interaction may pull S4-S5 linker towards TRP helix, resembling a gearbox-like motion. The S4-S5 linker undergoes a helix-to-loop transition, which may disrupt hydrophobic interactions between I604 (S4-S5 linker), I732 (TRP), and V725 (TRP-S6) (Figure 3c-d). The disruption of this interaction could potentially cause the N-terminus of the S5 helix to shift away from the S6 helix by ∼ 10 Å (Figure 3d). The decoupling motion between the S5 and S6 helices leads to the disruption of some interactions between the helices, such as those between D613 and T721, and I609 and V722 (Figure 3e). The increased space between the S5 and S6 helices allows the gating residue M718 to flip its side chain toward the pore.

A series of hTRPV4 mutants was generated to study gating-related interactions. They exhibited GSK101 responses comparable to the wild-type, confirming their suitability for further analysis (**Figure S8**). To test the importance of the proposed salt bridges interaction between R594, D743, and R746, as well as the hydrophobic interactions between I732, V725, and I604 for the TRPV4 inhibition by AH001, we evaluated the inhibitory activity of AH001 on hTRPV4 channels with mutants at R594A, D743A, D743E, and I732A. We found that compared to wild-type, the D743E mutation substantially decreased the potency of AH001 by 10-fold (Figure 3f), whereas the D743A mutant made the agonist used lose efficacy (**Table S3**). In addition, mutations of R594 and I732 to alanine decreased the inhibitory activity of AH001 (Figure 3f). These results suggested that the formation or disruption of the two interaction networks plays a critical role in the ligand-dependent gating of TRPV4. Furthermore, we hypothesized that the electrostatic interaction between K608 (S5) and E720 (S6) facilitates the rotation of the M718 gate. A substantial decrease in AH001 activity was marked by the E720A (IC_50_ = 38.56 μM) and K608A (IC_50_ = 14.66 μM) mutants (Figure 3f; **Table S3**, Supporting Information). These findings support our hypothesis that the decoupling of the S5 and S6 helices, along with the electrostatic interaction between K608 and E720 contribute to the clockwise rotation of the M718 gate, enabling the transition to the closed state.

In contrast to the single point mutations of K535A, I732A and E720A, as well as the double point mutation of K535A & E720A, which led to a partial reduction in the inhibitory activity of AH001 (Figure 3f; **Table S3**, Supporting Information), the triple point mutation K535A & I732A & E720A entirely eradicated the agonist’s ability to induce currents (**Table S3**, Supporting Information). The phenomenon was similarly observed in the triple mutation K535A & Q736A & E720A (**Table S3**, Supporting Information). Our computational and experimental data robustly support the potential existence of a signal transduction pathway during gating.

### 2.4 AH001 diminishes DRG neuronal hyperexcitability via TRPV4 interaction

To further verify the antagonistic activity of AH001 on the TRPV4 channel, we quantified the inhibitory effects of AH001 on TRPV4 in DRG neurons (Figure 4a-b). We established a chronic inflammatory pain model by using complete Freund’s adjuvant (CFA). Subsequently, electrophysiologic recordings were performed in small to medium-sized dorsal root ganglia (DRG) neurons from lumbar L4-L5 segments of mice^22, 23^, 3 days post-CFA induction, using a voltage ramp from −100 mV to +100 mV for the characterization of channel activity. Perfusion with 100 nM GSK101 induced outwardly rectifying TRPV4 currents, which were significantly attenuated by co-application of 20 μM AH001 (9.82 ± 4.33 pA/pF compared with 28.79 ± 13.78 pA/pF, *P* = 0.03, n = 6). Given the expression of TRPV4 in DRGs is closely implicated in nociception, we sought to examine whether AH001 contributes to antinociceptive effects by suppressing the hyperexcitability of peripheral sensory neurons. Lumbar L4-L5 DRG neurons from wild type or TRPV4 KO mice were isolated and treated with either the vehicle or AH001 (20 µM) for 2 h followed by patch-clamp recordings to assess excitability (**Figure S6a**, Supporting Information). No statistically significant differences were detected in the resting membrane potential of DRG neurons (**Figure S6b**, Supporting Information). Compared to the control group, neurons from mice in CFA group exhibited an increased action potential frequency and drop in the rheobase of nociceptors, indicating a pronounced hyperexcitable state (Figure 4c**, e-f**). The administration of AH001 (20 µM) considerably mitigated the nociceptor hyperexcitability induced by chronic inflammatory pain (Figure 4c**, e-f**). This reduction in hyperexcitability was evidenced by a decrease in action potential frequency (AH001 *vs.* vehicle = 1.46 ± 0.57 Hz *vs.* 6.55 ± 1.01 Hz, *P* = 0.0006) (Figure 4e) and an increase in rheobase (AH001 *vs.* vehicle = 42.9 ± 4.7 pA *vs.* 18.9 ± 2.7 pA, *P* = 0.002) (Figure 4f), with no discernible effect on the excitability of neurons from TRPV4 KO mice. In addition, AH001 did not affect the threshold potential of neurons, either in WT mice or in TRPV4 KO mice (Figure 4g). Collectively, these results indicate that AH001 effectively alleviates the aberrant hyperexcitability of DRG neurons induced by pain via TRPV4 interaction.

**Figure 4.**
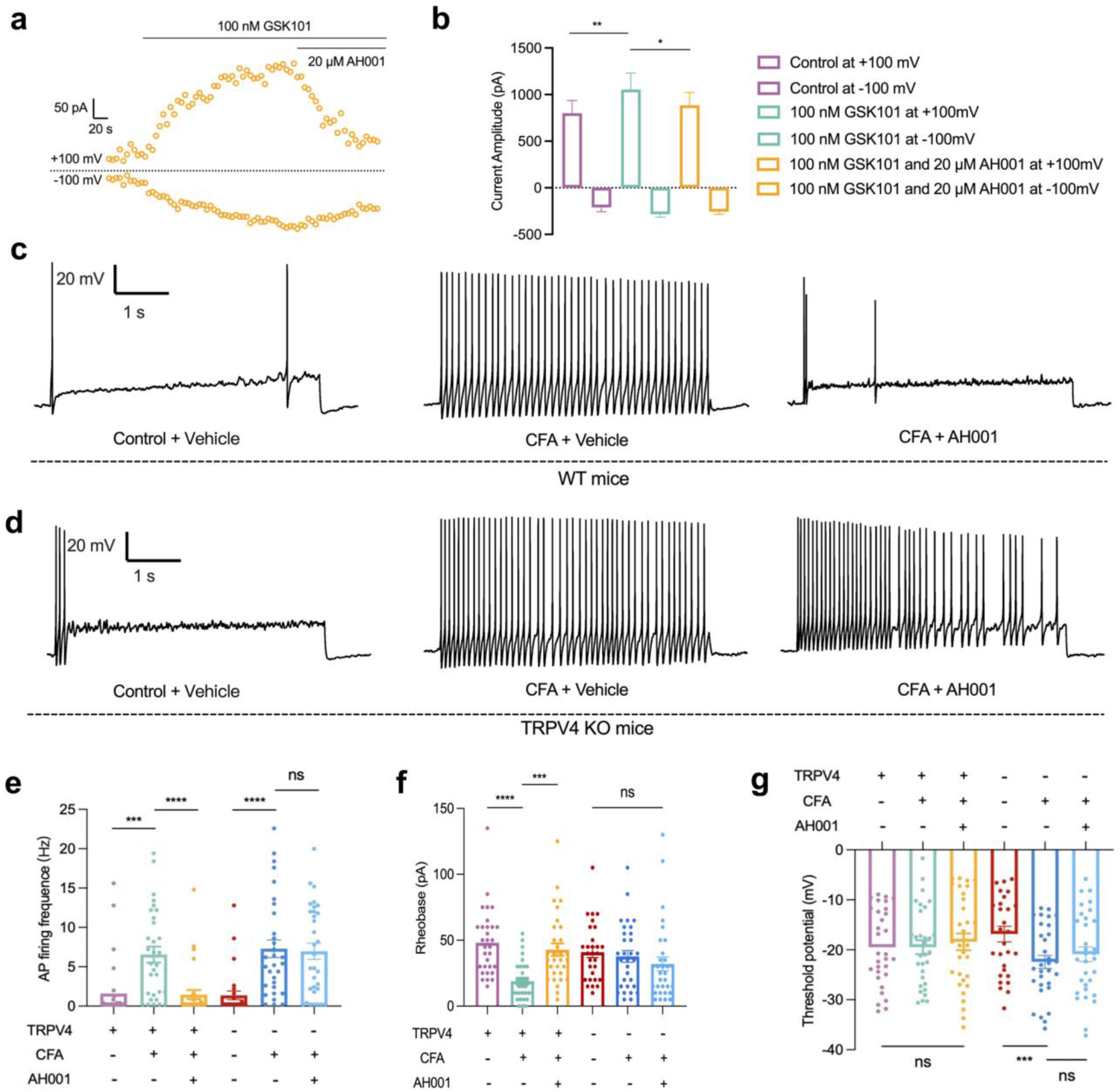
AH001 mitigates the hyperexcitability of dorsal root ganglion (DRG) neurons in CFA model mice by inhibiting TRPV4. **a**, Representative time course of whole-cell TRPV4 current obtained from voltage ramp (−100 mV to +100 mV every 5 seconds) in DRG neurons. Cells were held at 0 mV to inactivate voltage-gated calcium and sodium channels. **b**, Whole-cell recordings of peak current amplitude measured at +100 mV and −100 mV of DRG neurons. **c-d**, Action potential recordings from DRG neurons in control and CFA model mice, with or without AH001 treatment, and under conditions of TRPV4 knockout (**c**) or retention (**d**). **e-g,** Bar graph showing the effects of 20 μM AH001 on firing frequency (**e**), rheobase (**f**) and threshold potential (**g**) in nociceptive neurons, n = 30 cells in six mice, one-way ANOVA with Tukey’s post hoc test. \*\*\**P* < 0.001, and \*\*\*\**P* < 0.0001.

### 2.5 AH001 elicits analgesic effects in animal models through TRPV4

To evaluate the potential of AH001 as a TRPV4 inhibitor *in vivo*, we conducted a comprehensive assessment of its analgesic properties across various animal pain models. In the formalin-induced acute inflammatory pain model, AH001 exhibited significant anti-inflammatory analgesic activity, evidenced by a dose-dependent reduction in total licking time during the late phase of the model (Figure 5a). Encouraged by these results in the formalin model, we proceeded to test a 150 mg/kg dosage of AH001 in subsequent experiments to further investigate its efficacy in chronic and neuropathic pain models.

**Figure 5.**
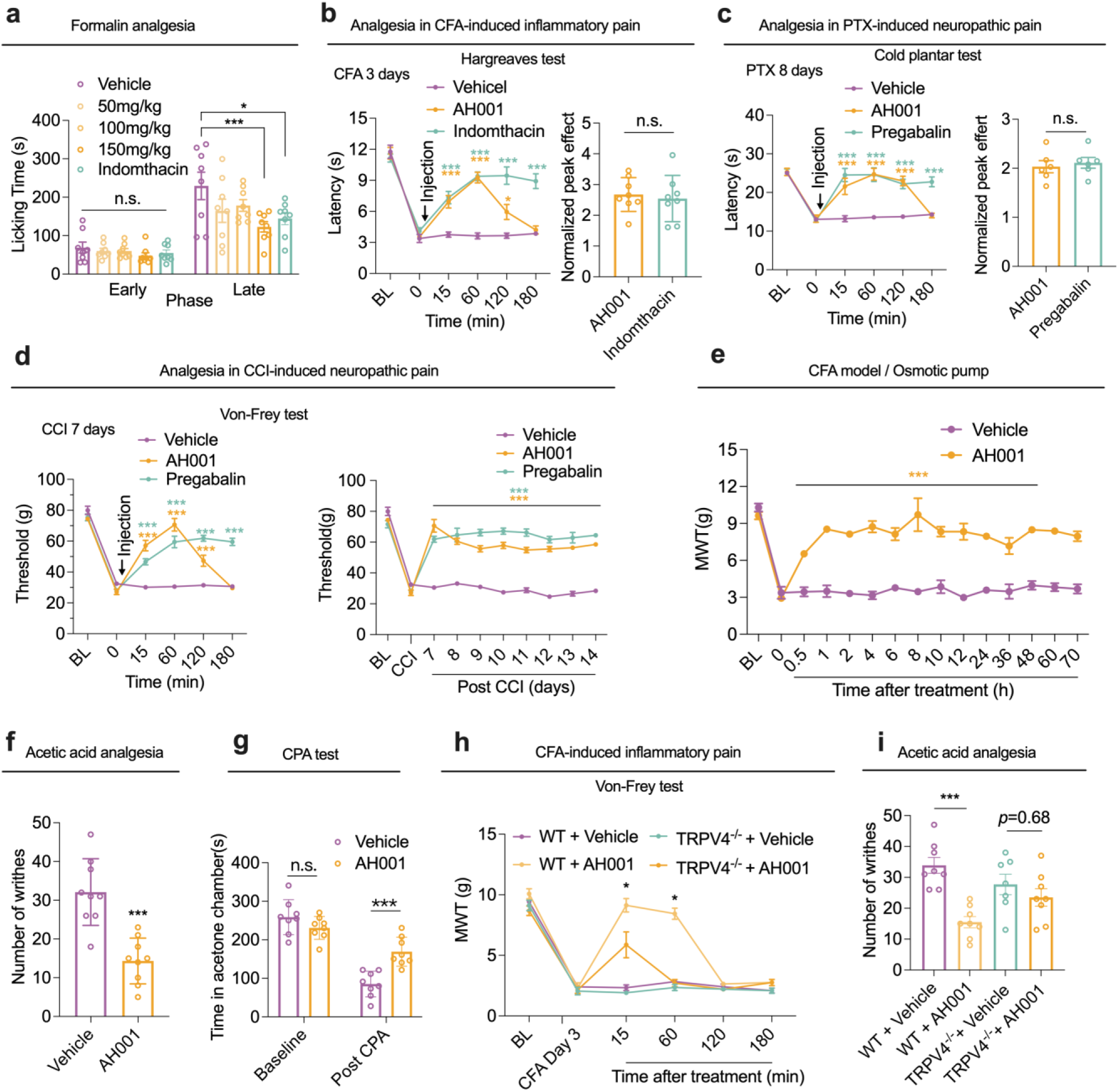
AH001 relieves pain in mouse models by inhibiting TRPV4. **a**, Dose-dependent analgesic effects of AH001 and indomethacin in formalin test. One-way ANOVA followed by Tukey post-tests with the vehicle-treated group, n = 8. **b,** Time course for analgesia by AH001 and indomethacin in Hargreaves test on day 3 after CFA-induced pain model (left). Two-way ANOVA compared with the vehicle-treated group, n = 8; Compared of normalized peek analgesic effects of AH001 and indomethacin (right). Unpaired t test, n = 8. **c,** Time course for analgesia by AH001 and pregabalin in cold plantar test on day 8 after PTX-induced pain model (left). Two-way ANOVA compared with the vehicle-treated group, n = 6; Compared of normalized peek analgesic effects of AH001 and pregabalin (right). Unpaired t test, n = 8. **d,** Time course for analgesia by AH001 and pregabalin in Von Frey test on day 7 and during days 7 to 14 post-CCI induction. Two-way ANOVA compared with the vehicle-treated group, n = 8. **e,** Long-term analgesic effect of AH001 by using subcutaneous implantable osmotic pump. MWT means mechanical withdrawal threshold. Two-way ANOVA compared with the vehicle-treated group, n = 3. **f,** Analgesic effects of AH001 in acetic acid-induced writhing test. Unpaired t test, n = 8. **g**, Quantification of two-chamber conditioned place aversion assay. Two-way ANOVA compared with the vehicle-treated group, n = 6. Data are shown as means ± SEM. \**P* < 0.05, \*\**P* < 0.01, and \*\*\**P* < 0.001. **h,** Time course for analgesia by AH001 in WT mice and TRPV4 KO mice on day 3 after CFA-induced pain model. Two-way ANOVA compared with the vehicle-treated group, n = 6. **i,** Analgesic effects of AH001 on WT mice and TRPV4 KO mice in acetic acid-induced writhing test. Two-way ANOVA compared with the vehicle-treated group, n = 8. Data are shown as means ± SEM. \**P* < 0.05, \*\**P* < 0.01, and \*\*\**P* < 0.001.

In the chronic inflammatory pain model induced by complete Freund’s adjuvant (CFA), AH001 effectively alleviated heat hyperalgesia in the Hargreaves test, achieving peak analgesic efficacy comparable to 10 mg/kg indomethacin (Normalized peek effect: AH001 *vs.* indomethacin: 2.68 ± 0.20 *vs.* 2.54 ± 0.27, *P* = 0.43) (Figure 5b). Similarly, in the paclitaxel (PTX)-induced neuropathic pain model, AH001 attenuated the cold hyperalgesia in the cold plantar test (Figure 5c) and mechanical allodynia in the Von Frey test (**Figure S7a**, Supporting Information). In a mouse model of chronic constriction injury of the sciatic nerve (CCI)–induced neuropathic pain, AH001 alleviated mechanosensitivity reactions during both the development phases (CCI 7 days, Figure 5d **left**) and maintenance phases (CCI 14 days, Figure 5d **right**) of pain, achieving peak analgesic efficacy comparable to pregabalin, a wildly used medicine for neuropathic pain treatment (**Figure S7b**, Supporting Information). Moreover, AH001 effectively alleviated heat hyperalgesia in CCI model (**Figure S7c**, Supporting Information). While the analgesic effects of AH001 were transient, likely due to its rapid metabolism *in vivo* (**Table S4**, Supporting Information), we addressed this issue by investigating sustained administration through a subcutaneous implantable osmotic pump. This approach provided stable long-term analgesia without significant side effects (*P* < 0.001) (Figure 5e). Remarkably, AH001 also demonstrated potential effectiveness against visceral pain, as it significantly decreased the number of writhing movements of animals during the acetic acid-induced writhing test (writhing times: AH001 *vs.* indomethacin = 14.3 ± 2.9 *vs.* 32.1 ± 2.0, *P* < 0.001) (Figure 5f).

To further explore the analgesic potential of AH001, we employed a two-chamber conditioned place aversion (CPA) assay in mice with PTX-induced neuropathic pain. Following pre-conditioning, mice with PTX-induced neuropathic pain were administrated AH001 or the vehicle 1 h prior to testing. As long as the mice remained in the right chamber, we repeatedly stimulated their left hind paw with 25 µL of acetone (**Figure S7d**, Supporting Information). Compared to the baseline phase, the vehicle group spent less time in the right chamber, suggesting the occurrence of aversion-like behavior. In contrast, administration of AH001 significantly attenuated this aversion (Figure 5g).

To confirm that the analgesic effects of AH001 are mediated by TRPV4 inhibition, we utilized TRPV4 knockout (TRPV4^-/-^) mice. In the CFA-induced chronic inflammatory pain model, AH001 administration (150 mg/kg) significantly increased hind paw withdrawal thresholds in wild-type (WT) mice but had no effect in TRPV4^-/-^ mice (Figure 5h). Furthermore, in the acetic acid-induced writhing test, AH001 failed to reduce visceral pain in TRPV4^-/-^ mice (Figure 5i). These findings conclusively demonstrate that the analgesic effects of AH001 are dependent on TRPV4, providing robust evidence for its mechanism of action.

## 3. Discussion

In this study, we identified AH001, a natural product, as a selective and potent inhibitor of TRPV4 using whole-cell patch-clamp recordings. Mechanistically, our findings expand upon prior studies that have emphasized the critical roles of the S4–S5 linker and TRP helix in TRP channel gating transitions^11, 24, 25^. Specifically, the gearbox-like model has postulated that TRP channel gating is regulated by coupling between these regions^26^. Additionally, Lee et al.^15^ hypothesized that antagonists promote TRPV4 channel closure by decoupling the interaction network among the VSLD (primarily the S2-S3 helices), the TRP helix, and CD domain. However, prior to this work, the detailed molecular mechanisms underlying antagonist-induced conformational changes—including the sequence of binding interactions, coupling events within the gating machinery, and the final channel closure—remained elusive. Combining structural analysis and mutagenesis studies, our study proposes the potential gating mechanism of TRPV4, advancing the current understanding of TRP channel modulation. In brief, AH001 was found to bind specifically within the VSLD cavity, potentially triggering a decoupling motion between the S2-S3 helices and the TRP helix. The S4-S5 linker is pulled towards the TRP helix, resembling a gearbox-like motion. The remodeling of the S4-S5 linker may induce decoupling between the S5 and S6 helices, ultimately resulting in a clockwise rotation of M718 and gate closure. This cascade may further be stabilized by electrostatic interactions between K608 and E720, underscoring the intricate network of molecular forces governing TRPV4 inhibition (Figure 6).

**Figure 6.**
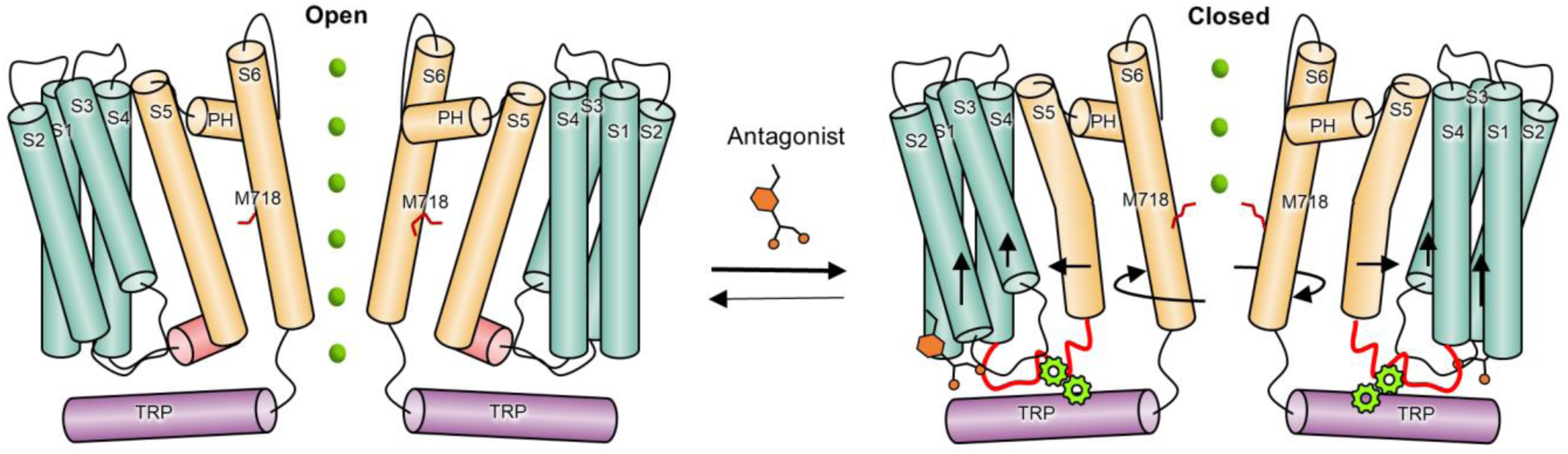
Schematic representation of the proposed potential gating mechanism of TRPV4 inhibition by AH001. We propose a three-step model to elucidate the conformational transition of TRPV4 from an open to a closed state upon the binding of AH001: (1) AH001 binds within the VSLD cavity, potentially triggering a decoupling motion between the S2-S3 helices and the TRP helix, (2) the S4-S5 linker is pulled towards the TRP helix, resembling a gearbox-like motion, (3) the remodeling of the S4-S5 linker may induce decoupling between the S5 and S6 helices, ultimately resulting in a clockwise rotation of M718 and gate closure. Black arrows denote potential movements upon ligand binding.

Importantly, our study also demonstrates that AHP, the glucuronide metabolite of AH001, retains potent antagonist activity against TRPV4. The cryo-EM structure of TRPV4 in complex with AHP revealed its binding site within the VSLD cavity, highlighting the structural similarities and differences between AH001 and AHP. This structural characterization offers valuable insights for optimizing AH001 and developing related analogues with enhanced efficacy and pharmacokinetic properties. The identification of an active metabolite further underscores the clinical potential of AH001, as metabolites often contribute significantly to a compound’s *in vivo* pharmacological profile.

Beyond mechanistic insights, our findings highlight the therapeutic applicability of AH001 as a promising candidate for pain management. AH001 demonstrated robust efficacy in alleviating pain in cellular and animal models by selectively inhibiting TRPV4 activity. This selectivity is particularly significant, as it minimizes off-target effects on other TRP channels, a common challenge in TRP channel-targeted drug discovery. Notably, AHP, the major glucuronide metabolite of AH001, also retained inhibitory activity against TRPV4. The ability to precisely target TRPV4 without affecting closely related channels positions AH001 and AHP as promising leads for analgesic development.

Collectively, this study represents an advance in TRP channel biology and pharmacology. By uncovering the molecular basis of TRPV4 inhibition, characterizing a natural product and its active metabolite, and demonstrating their therapeutic potential, our work bridges fundamental mechanistic insights with translational applications. These findings not only provide a strong foundation for the development of selective TRPV4-targeted analgesics but also pave the way for innovative strategies to tackle TRPV4-associated diseases, such as neuropathic pain, osteoarthritis, and inflammatory disorders. This work exemplifies the potential of integrating structural biology, pharmacology, and drug discovery to address unmet medical needs.

## 4. Experimental Methods

### 4.1 Electrophysiology

AH001 was prepared to 1M stock solution in water. Before adding into the external solution, AH001 solutions were shaken for 20 min and warmed to 37 °C. Then, the stock solution was diluted into the working solution successively with the extracellular solution. All concentrations dissolve without visible precipitation.

Electrophysiological tests of hTRPV1, hTRPA1 and hTRPM8 were performed with stable cell lines. These stable cell lines were based on a standard HEK293 cell line, which were provided by ICE Bioscience (Beijing, China). Electrophysiology tests of hTRPV3, hTRPV4 and mutations of hTRPV4 were performed with transiently transfected HEK293 cells. The cDNAs of hTRPV3 (NM_001258205.2) and hTRPV4 (NM_021625.5) were subcloned into the pcDNA3.1 vector (Invitrogen). All hTRPV4 mutants were generated by site-directed mutagenesis. All mutant plasmids used in this study were verified by DNA sequencing.

Whole-cell patch-clamp recordings were performed at room temperature using a HEKA EPC10 amplifier with PatchMaster software (HEKA). Currents were filtered at 2 kHz and sampled at 10 kHz. For ramp recordings, the pipette solutions contained 50 mM CsCl, 10 mM NaCl, 10 mM HEPES, 60 mM CsF and 20 mM EGTA (buffered to pH 7.2 with CsOH), and bath solutions contained 140 mM NaCl, 3.5 mM KCl, 1 mM MgCl_2_•6H_2_O, 2 mM CaCl_2_•2H_2_O, 10 mM D-Glucose, 10 mM HEPES, 1.25 mM NaH_2_PO_4_•2H_2_O (buffered to pH 7.4 with NaOH). Cells were held at 0 mV before given a ramp stimulus from −100 mV to +100 mV every 5 s. The dose-response curves were fitted with the Hill equation: Y=1/(1+10^((LogIC_50_-X) *HillSlope)), IC50 calculation and curve fitting were performed using GraphPad Prism software. All recordings were performed at room temperature and all drugs were applied to the bath by gravity at the same rate. All agonists were purchased from Sigma-Aldrich.

### 4.2 Protein expression and purification of TRPV4

The human TRPV4 construct (amino acids 148 to 787) was cloned into the pEGBacMam vector. The resulting protein contains 8 × His-Flag tag on its carboxyl terminus. P4 baculovirus was produced in the Bac-to-Bac Baculovirus Expression System (Invitrogen) using Spodoptera frugiperda (Sf9) cells. HEK293F [from the American Type Culture Collection (ATCC)] cells were infected with 10% (v/v) P4 baculovirus at a density of 2.5 × 10^6^ cells/ml for protein expression at 37°C. After 12 h to 18 h, 10 mM sodium butyrate was added, and the temperature was reduced to 30°C. Cells were harvested 72 h after transduction and frozen at −80°C.

Before solubilization, cells were resuspended in a buffer containing 20 mM Tris, pH 8.0, 0.0025 mg/mL Leupeptin (BIORIGIN), 1 mM PMSF (Beyotime) with 10 U/mL Benzonase Nuclease (HaiGene), and cell membranes were disrupted by dounce homogenization. Lysed membranes were collected by ultracentrifugation for 60 min at 15,000 rpm and the precipitate was re-suspended and homogenized by dounce in a buffer containing 1.0% (wt/vol) Lauryl maltose neopentyl glycol (LMNG; Anatrace), 0.1% (wt/vol) cholesteryl hemisuccinate (CHS; Anatrace), 20 mM Tris, pH 8.0, 150 mM NaCl and 0.0025 mg/mL Leupeptin. Solubilization proceeded for 2.5 h at 4 °C, followed by ultracentrifugation for 60 min at 41,000 rpm. After centrifugation, the solubilized supernatant was incubated with Anti-Flag resin for 2 h at 4 °C. The resin was washed with 30 column volumes of 20 mM Tris, pH 8.0, 150 mM NaCl, 0.03% LMNG, 0.003% CHS and 0.0025 mg/mL Leupeptin. The protein was eluted with 10 column volumes of 20 mM Tris, pH 8.0, 150 mM NaCl, 0.03% LMNG, 0.003% CHS, 0.0025 mg/mL Leupeptin and 0.3 mg/mL Flag peptide. Finally, the protein was then concentrated to 1.0 mL with a 50-kDa molecular weight cut-off concentrator (Millipore) before further purification on Superose 6 size exclusion column (Cytiva) in 20 mM Tris-HCl pH 8.0, 150 mM NaCl, 1 mM DTT, 0.01% GDN (Anatrace), 0.0025 mg/mL Leupeptin. The peak fractions were collected and concentrated to 5 mg/mL for Cryo-EM sample preparation.

### 4.3 Cryo-EM sample preparation and data collection

Purified hTRPV4 protein at the concentration of 5-8 mg/mL was incubated with AHP at a 1:20 molar ratio on ice for 40 min for the next step centrifugation (16200 g, 4℃ for 5 min). The TRPV4-AHP complex sample was then applied for cryo-EM grid preparation.

Cryo-EM micrographs were collected on a 300 kV Thermo Fisher Titan Krios G4 electron microscope equipped with a Falcon4 direct detection camera. The micrographs were collected at a calibrated magnification of x96,000, yielding a pixel size of 0.404 Å at a super-resolution mode. In total, 17,269 micrographs were collected at an accumulated electron dose of 50 e^-^Å^-2^ on each micrograph that was fractionated into a stack of 32 frames with a defocus range of −1.0 μm to −2.0 μm. The exposure time and dose rate were 2.6 s and ∼19 e^-^pixel^-1^s^-1^.

### 4.4 Image processing and 3D reconstruction

Beam-induced motion correction was performed on the stack of frames using MotionCorr2^27^. The contrast transfer function (CTF) parameters were determined by CTFFIND4^28^. A total 17,269 micrographs were collected and the data were processed using CryoSPARC^29^. The data processing workflow is summarized in **Figure S3**. All model were built by fitting a structure (predicted by AlphaFold2) into the density map using UCSF Chimera^30, 31^, followed by a manual model building of the complex molecules in COOT^32^ and a real space refinement in PHENIX^33^. The geometry statistics for models were generated using MolProbity and the model statistics were listed in **Table S1**.

### 4.5 Safety screen study

For evaluation of off-target effects, AH001 at 10 μM in duplicate was screened for the 44 safety panel following the manufacturer’s instructions with a commercial screen (SafetyScreen 44, ICE Bioscience). The experiment was repeated at least twice. The percent activation or inhibition of the positive control ligands specific for each target was used to calculate the binding. The percent inhibition of control enzyme activity was used to calculate the compound’s enzyme inhibition effect. The safety screening results are summarized in **Table S2** and Figure 2k.

### 4.6 Animals

All experimental protocols were approved by the Institutional Animal Care and Use Committee of Sichuan University West China Hospital (protocol No.: 20220127001). Animal experiments were conducted in strict accordance with the National Institutes of Health Guide for the Care and Use of Laboratory Animals. All wild-type C57BL/6 mice were purchased from Dashuo experimental animal corporation (Chengdu, Sichuan, China). Trpv4 knock-out mice (Trpv4^-/-^) were generated by Cyagen Biosciences (Suzhou, Jiangsu, China). The animals had unrestricted access to clean water and food and were housed in a conventional facility at 22 ± 2 °C on a 12-hour light/dark cycle.

#### 4.6.1 TRPV4 KO mice generation

In order to generate a TRPV4 knockout C57BL/6N mouse line with the CRISPR-Cas9 genome editing system, two single-guide RNAs (sgRNA-1, 5′-CAGGTGGTCGAGTACCAGCCCGG-3′, and sgRNA-2, 5′-CTTGCATAGTAGGGTGCTAGGGG-3′) were designed (Cyagen Biosciences). A 5703-bp deletion was bound from exon 3 to exon 5 of the TRPV4 gene locus, resulting in TRPV4 KO mice with a frameshift mutation.

#### 4.6.2 Formalin test

The mice were placed in plexiglas chambers individually to adapt for at least 30 minutes. 5% formalin (Sigma-Aldrich, St Louis, Missouri, USA) was dissolved in saline. 45 minutes after intragastric administration of vehicle or AH001, 20 µL formalin was subcutaneously injected into the left hind paw of mice. And then, the mice were placed back into plexiglas chambers, where movement of the mice was recorded by an action camera. The number of paw flinches was counted within the time phase I (0–5 min) and phase II (15–30 min).

#### 4.6.3 Chronic inflammatory pain model

20 μL Complete Freund’s Adjuvant (CFA) (Sigma-Aldrich) was injected into the left hindpaw of the mice subcutaneously to induce chronic inflammatory pain in mice. Behavioral test was carried out after three days of injection.

#### 4.6.4 Paclitaxel injection model

Mice were intraperitoneally injected with paclitaxel solution 2 mg/kg every other day. After 4 times of paclitaxel administration, behavioral test was carried out to measure the pain threshold of mice. Paclitaxel (Yangzi Pharmaceutical Co., Ltd. Taizhou, Jiangsu, China) was diluted with a solvent (ethanol: hydrogenated castor oil: normal saline = 4:4:17).

#### 4.6.5 Chronic constriction injury model

Under anesthesia inhaled by isoflurane, the left sciatic nerve of mice was exposed. After the sciatic nerve trunk was isolated, two ligations were made with chrome catgut, and the tightness was suitable for the slight twitch of the left hind limb of mice during ligation. 7 days after the surgery, behavioral test was conducted.

#### 4.6.6 Acetic acid-induced writhing test

Briefly, the mice were treated with vehicle or AH001. After 45 min, they were injected i.p. with 1% acetic acid (0.01 mL/g), and the stretching episodes were recorded for 20 min.

#### 4.6.7 Behavior tests Cold plantar test

The mice were placed in an elevated transparent cage (20 × 20 × 14 cm) 2-3 h daily for three days. During the test, the dry ice was pointed at the left rear paw of the mice, and the time of shrinkage/licking/shaking of the mice was recorded. The test was repeated three times at an interval of 5 minutes, and the average value was recorded. To avoid tissue damage, the cut-off time is set at 35 s.

##### von Frey test

The mice adapted to the same conditions as before. After acclimation, an electronic von Frey tester (no. 2391, IITC Inc. Woodland Hills, CA) was used for mechanical paw withdrawal threshold measurement. If rapid foot shrinking/licking response occurred during the electronic von Frey test, the value was recorded as positive, and the average value is recorded three times.

##### Hargreaves test

The mice adapted to the same conditions as before. For thermal withdrawal latency, the measurements were made using a classic infrared radiometer (Ugo-Basile, Italy) with a temperature set at 35°C. During the test, the radiation light source was pointed at the right rear paw of the mice, and the incubation period of foot contraction/licking in the mice was taken as thermal withdrawal latency (TWL). The test was repeated three times at a 5-minute interval, and the average value was recorded. To avoid tissue damage, the cut-off time is set at 20 seconds.

##### Conditioned place aversion test

A three-chamber apparatus with walls of different colors and floors of different textures was used. The three chambers were referred to as the start chamber, acetone chamber and contralateral chamber. During the pre-stimulation stage, mice were allowed to explore the three chambers for 10 min. An hour after receiving AH001 (150 mg/kg, i.g.) or vehicle, each mouse was imprisoned to an acetone chamber and exposed to 50 μL acetone splashing for 10 min (stimulation stage). Following the stimulation stage, mice were allowed to move freely again among the three chambers for 10 min (post-stimulation stage). The time that each mouse spent in each chamber was recorded by the tracking system Smart v2.5 (Panlab, Barcelona, Catalunya, Spain).

### 4.7 DRG neurons preparation and electrophysiology

Mice on the third day after CFA injection were sacrificed and the DRG neurons were isolated. The DRGs were collected in a 35-mm tissue culture dish with HBSS (Hank’s Balanced Salt Solution) and the nerve fibers were cut. Then, the DRGs were digested by 2 mL collagenase IV (2 mg/mL) and 2 mL 1% trypsin for 30 min at 37 ℃, respectively. The digestion was stopped by 200 μL fetal bovine serum and the ganglia gently washed with a neurobasal medium (Gibco). Next, the neurons were seeded on polyornithine and laminin-coated glass coverslips. Neurons were fed with 2 mL neurobasal medium supplemented with 2% B27 (Gibco), 1% GlutamaxTM (Gibco) and cultured at 37 ℃ in 5% CO_2_ for 2-4 h. For recording the action potential, the pipette solution contained (in mM): 120 KCl, 4 NaCl, 1 MgCl_2_, 0.5 CaCl_2_, 10 EGTA, 10 HEPES, 2 Na_2_-ATP (buffered to pH 7.2 with KOH). The bath solution contained (in mM): 130 NaCl, 3 KCl, 1 CaCl_2_, 2 MgCl_2_, 1 NaH_2_PO_4_, 10 HEPES, 10 glucose (buffered to pH 7.3 with NaOH). Neurons were current-clamped via the whole cell configuration of the patch clamp with an Axopatch 700B amplifier (Molecular Devices, Sunnyvale, CA, USA), DigiDATA and pClamp software. The micropipettes for whole-cell recordings were pulled from Flaming Brown micropipette puller (P-1000, Sutter Instrument, Novato, CA) to obtain electrode resistances ranging from 2.0 to 4.0 MΩ. The digitized current was sampled at 10 kHz and filtered at 2 kHz for analysis. Current threshold was determined by a series of depolarizing currents from 0 to 200 pA in 5-pA step increments. The threshold current triggering the action potential was recorded as the rheobase. Firing frequency of action potentials was measured in 5 s at 2-fold rheobase injection.

### 4.8 Statistical analysis

Curve fitting analyses and statistical analyses of behavior experiments were performed using GraphPad 9.5.1 and presented as mean ± SEM. The level of statistical significance was set at *P* < 0.05. Two-tailed paired Student’s t-tests and one-way or two-way ANOVA followed by Tukey post tests were used to examine statistical significance: n.s. indicates no significance. *, ** and *** indicate *P* values < 0.05, < 0.01, and < 0.001, respectively.

## Acknowledgments

The authors gratefully acknowledge financial support from National Natural Science Foundation of China (82425104 to H.L., 82273784 to B.K.); the National Key R&D Program of China (2022YFC3400501, 2022YFC3400504); the 1.3.5 Project for Disciplines of Excellence, West China Hospital, Sichuan University (ZYYC21002); S.L. is also sponsored by the Shanghai Rising-Star Program (23QA1402800).

## Author contributions

H.L. and B.K. initiated the project. H.L. and B.K. conceived and codirected the study; S.L., H.L. and B.K. discovered the antagonist AH001 inhibiting TRPV4, and S.L., H.L. and B.K. coordinated the cryo-EM related studies; Z.Y.,S.R. and S.L. performed the simulation and analysis of cryo-EM data; J.Q., H.Z., Y.G., Z.L. and B.K. performed electrophysiology, DRGs and *in vivo* animal studies; J.W. and R.W. was in charge of the *in vitro* pharmacological profiling study; S.L. and L.Z. helped the cryo-EM study; N.C., T.Y., W.L. and Z.Z. synthesized the compounds; Y.D. helped the pharmacokinetics study; Y.Z., R.W., Y.G. and J.L. analyzed and dealt with the animal data; H.L. and B.K. interpreted and reviewed all data; Z.Y., J.Q., S.L. and Y.G. wrote the manuscript; H.L. and B.K. revised the article.

## Data Availability Statement

Atomic coordinates and cryo-EM map of the reported structure have been deposited into the Protein Data Bank (PDB) and Electron Microscopy Data Bank (EMD) under the session codes PDB 9JPD and EMD-61697 for TRPV4 in complex with AHP.

